# Monitoring contractility in single cardiomyocytes and whole hearts with bio-integrated microlasers

**DOI:** 10.1101/605444

**Authors:** Marcel Schubert, Lewis Woolfson, Isla RM Barnard, Andrew Morton, Becky Casement, Gavin B Robertson, Gareth B Miles, Samantha J Pitt, Carl S Tucker, Malte C Gather

## Abstract

Cardiac regeneration and stem cell therapies depend critically on the ability to locally resolve the contractile properties of heart tissue^1,2^. Current regeneration approaches explore the growth of cardiac tissue *in vitro* and the injection of stem cell-derived cardiomyocytes^3–6^ (CMs) but scientists struggle with low engraftment rates and marginal mechanical improvements, leaving the estimated 26 million patients suffering from heart failure worldwide without effective therapy^7–9^. One impediment to further progress is the limited ability to functionally monitor injected cells as currently available techniques and probes lack speed and sensitivity as well as single cell specificity. Here, we introduce microscopic whispering gallery mode (WGM) lasers into beating cardiomyocytes to realize all-optical recording of transient cardiac contraction profiles with cellular resolution. The brilliant emission and high spectral sensitivity of microlasers to local changes in refractive index enable long-term tracking of individual cardiac cells, monitoring of drug administration, and accurate measurements of organ scale contractility in live zebrafish. Our study reveals changes in sarcomeric protein density as underlying factor to cardiac contraction which is of fundamental importance for understanding the mechano-biology of cardiac muscle activation. The ability to non-invasively assess functional properties of transplanted cells and engineered cardiac tissue will stimulate the development of novel translational approaches and the *in vivo* monitoring of physiological parameters more broadly. Likewise, the use of implanted microlasers as cardiac sensors is poised to inspire the adaptation of the most advanced optical tools known to the microresonator community, like quantum-enhanced single-molecule biosensing or frequency comb spectroscopy^10^.

---

To elucidate CM contractility under various experimental conditions, we explored the integration of WGM microlasers as multifunctional optical sensors. Chip-based fibre- and prism-coupled WGM biosensors have previously achieved sensitivities down to the single molecule and protein level^11,12^. However, their potential for intracellular sensing remains largely unexplored as integration into biological systems requires further miniaturization, self-sustained and prolonged emission of light, and data analysis protocols with improved robustness. Recently, microlasers were proposed as novel optical tags to uniquely discriminate hundreds of thousands of cells^13–16^.

Fig. 1 illustrates the general principle of our laser-based contraction sensor. Brightly fluorescent polystyrene microspheres with a diameter between 10 to 20 μm were used as efficient and robust microscopic WGM lasers that show multi-mode emission under remote optical pumping.^13^ These lasers were actively internalized by different types of cardiac cells (Supplementary Figs. 1 and 2). Upon CM contraction, individual peaks in the emission spectrum of the lasers showed a spectral red-shift (typically, Δλ ≈ 50 pm; Fig. 1d). Due to the bright and narrowband laser emission, the wavelength of each lasing mode can be monitored rapidly (acquisition rate, 100 Hz) and accurately (spectral resolution, 1 pm), revealing pulse-shaped perturbations in lasing wavelength synchronized across all modes and coincident with the spontaneous contractions of the cell (Figs. 2a, 2b and Supplementary Video 1). By tracking at least 2 pairs of TE and TM lasing modes and fitting their position to an optical model, we independently determined the diameter *D* of each microsphere and the average external refractive index *n*_ext_, i.e. the refractive index (RI) of the volume probed by the evanescent component of the WGM (Fig. 2c and Supplementary information). This revealed a characteristic increase in RI during cell contractions. Statistical analysis of the microsphere diameter was then applied to reduce the effect of fitting noise before reiterating the RI calculation. This significantly improved the signal quality and thus allowed the detection of minute changes in *n*_ext_, with a RI resolution of 5 × 10^−5^, which rivals the most sensitive cell refractometric techniques currently available^17^.

**Fig. 1.**
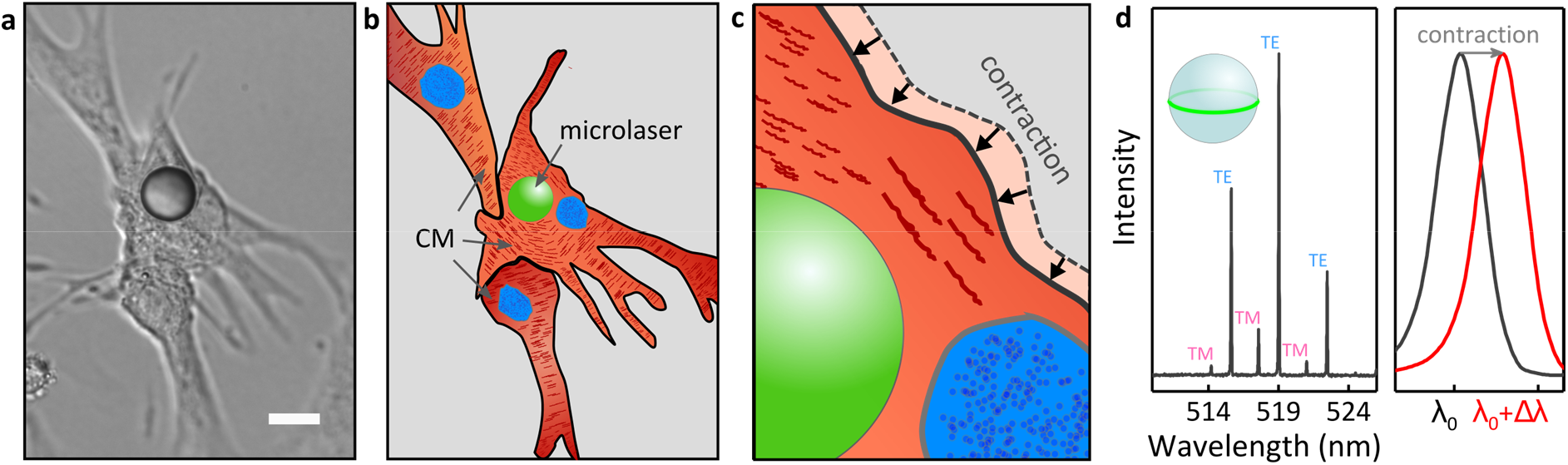
Principle of microlaser-based intracellular sensing in neonatal mouse CMs. **a**, DIC microscopy and **b**, schematic illustration of a group of neonatal CMs and an intracellular microlaser (green sphere). **c**, Magnified view visualizing the contractile movement of the cell around the microlaser, due to the action of sarcomeres (dark red fibres). **d**, WGM spectrum of a microlaser showing multi-mode lasing in pairs of TE- and TM-modes (left). WGMs are localized within an equatorial plane close to the surface of the microlaser (inset, green line). Zoom-in onto one peak in the WGM spectrum illustrating the red-shift in lasing wavelength upon CM contraction (right; λ_0_ = 519 nm, Δλ = 50 pm). Scale bar, 15 μm.

**Fig. 2.**
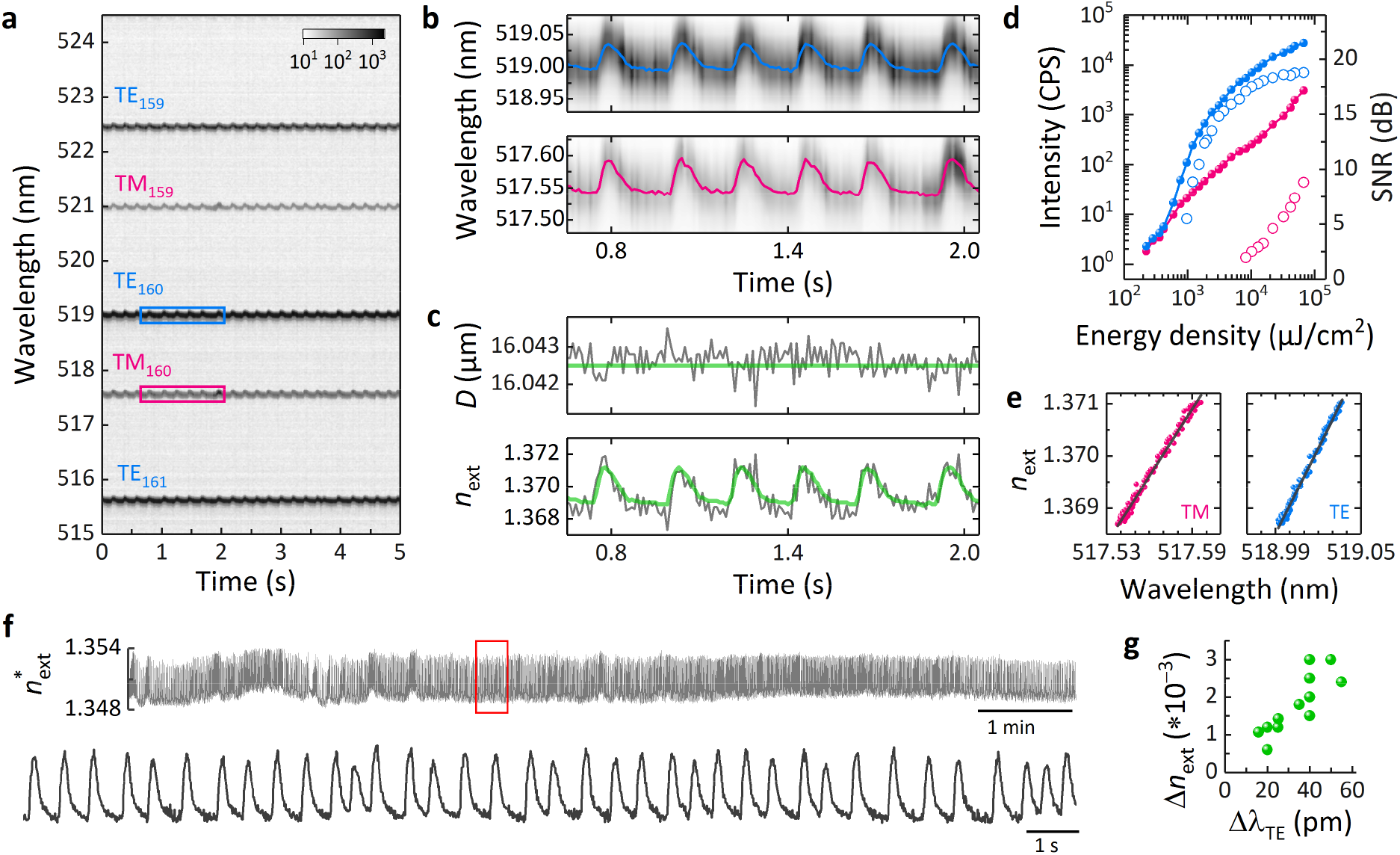
Transient red-shifts of microlaser emission are caused by changes in intracellular refractive index. **a**, Contour plot of the temporal evolution of the lasing spectra for an intracellular microlaser, measured with 10 ms temporal resolution. **b**, Magnified view of the areas highlighted in **a**, for a pair of TM (pink) and TE (blue) WGMs. The coloured lines show the centre position of each mode obtained from peak fitting. Shifts to longer wavelengths coincide with spontaneous CM contractions. **c**, Calculated diameter (top) of the microlaser (grey) and time-averaged diameter (green). External refractive index *n*_ext_ (bottom) calculated with unrestricted microlaser size (grey) and by applying the fixed mean diameter of the microlaser (green). **d**, Typical threshold characteristics (left axis, closed symbols) for the brightest TE mode (blue) and the least intense TM mode (pink) of 4 tracked lasing modes. Lasing thresholds are about 500 μJ/cm^2^ (TE) and 20 mJ/cm^2^ (TM), respectively. Signal-to-noise ratio (SNR) (right axis, open symbols) of the same modes under single pulse excitation. **e**, Mode calibration of the 2 modes shown in **b** using data from 6 contractions. From the slope of the linear fit (grey line), a sensitivity *S* of 0.0429 nm^−1^ and 0.0549 nm^−1^ is obtained for the TM and TE mode, respectively. **f**, Continuous single cell monitoring over 10 min (top) at 2 mJ/cm^2^ (corresponding to 2 nJ/pulse) and magnified view of the 20 s window indicated by the red rectangle (bottom). **g**, Average refractive index change (Δ*n*_ext_) between resting phase (diastole) and peak contraction (systole) for n=12 cells plotted over the corresponding average change of the dominant TE WGM (Δ*λ*_TE_).

Of the 2 pairs of TE and TM lasing modes required for fitting to the optical model, the brightest mode typically has a lasing threshold below 1 mJ/cm^2^ (corresponding to <1 nJ/pulse, Fig. 2d). Above threshold, this mode rapidly increased in intensity to become 2 to 3 orders of magnitude more intense than the bulk fluorescence of the microlaser. Single pulse excitation at around 1 mJ/cm^2^ can be therefore used to accurately determine the spectral position of this mode (Supplementary Fig. 3). By comparison, the least intense mode of the 2 pairs required 10 to 50 times higher pump energy to pass the lasing threshold and to determine its spectral position with sufficient accuracy to ensure convergence of our fitting algorithm. Furthermore, we found that the periodic changes in RI due to cardiomyocyte contraction can be utilized to determine the sensitivity (*S*) of each laser mode (Fig. 2e, Supplementary Fig. 4). Using *S* and tracking the spectral position of just the brightest lasing mode then allows calculation of a linearly approximated external refractive index 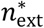, which means the pump energy can be reduced by at least one order of magnitude. This calibration protocol enabled real time RI sensing, allowed continuous yet non-disruptive read-out (Fig. 2f) and greatly improved the robustness of the approach under challenging experimental conditions (see below).

Analysis of multiple CMs furthermore revealed that contractions consistently led to an increase of cellular RI which indicates the presence of a highly reproducible physiological process that alters the optical properties of CMs depending on the activation state of their contractile elements (Fig. 2g).

To identify the origin of the RI increase during CM contraction we analysed the 3D organization of myofibrils, cellular organelles which comprise repeating contractile elements called sarcomeres. It is generally assumed that CMs contract under isovolumetric conditions^18^, yet X-ray diffraction experiments have revealed a linear relationship between sarcomere length and volume of the myofibril unit cell indicating that cell contractions significantly increase the protein density of the myofibrils^19^. 3D reconstructions of cells showed that microlasers are surrounded by and in direct contact with a dense network of myofibrils (Figs. 3a and 3b, Supplementary Fig. 5), indicating a strong overlap of the contractile protein machinery with the evanescent field of the laser mode, which typically extends up to 200 nm above the resonator surface. Cellular contractility in neonatal CMs was then measured by staining sarcomeric actin filaments and tracking their length change during the contraction cycle, while simultaneously recording spectral shifts in microlaser emission (Figs. 3c and 3d, and Supplementary Video 2). We find that the sarcomere length (SL) of myofibrils was directly correlated with *n*_ext_, indicating that structural changes inside myofibrils cause the red-shifts in lasing wavelength (Fig. 3b). Given that during contraction *n*_ext_ increased by up to 0.003 and using the known protein refractive index increment (*dn*/*dc*), we further estimated that the observed contraction-induced changes in sarcomere length by about 10% led to a maximum increase in protein concentration of about 8% (Supplementary information). This finding is consistent with the previously reported decrease in unit cell volume^19^. It does not contradict observations that the contraction of the whole heart is isovolumetric; during contraction, cardiac cells are likely to expel water from the myofibrils into different parts of the cell or to the extracellular space^20^, causing a local increase in myofibril density while still conserving the overall tissue volume. The effects associated with sarcomeric lattice spacing and unit cell changes are of great importance for the function of cardiac cells and are believed to play an important role in regulating the length-dependent activation of the heart (Frank-Starling law). Since transient RI profiles provide a direct measure of CM contractility and myofibril density, they provide new insights to the mechano-biology of cardiac cells.

**Fig. 3.**
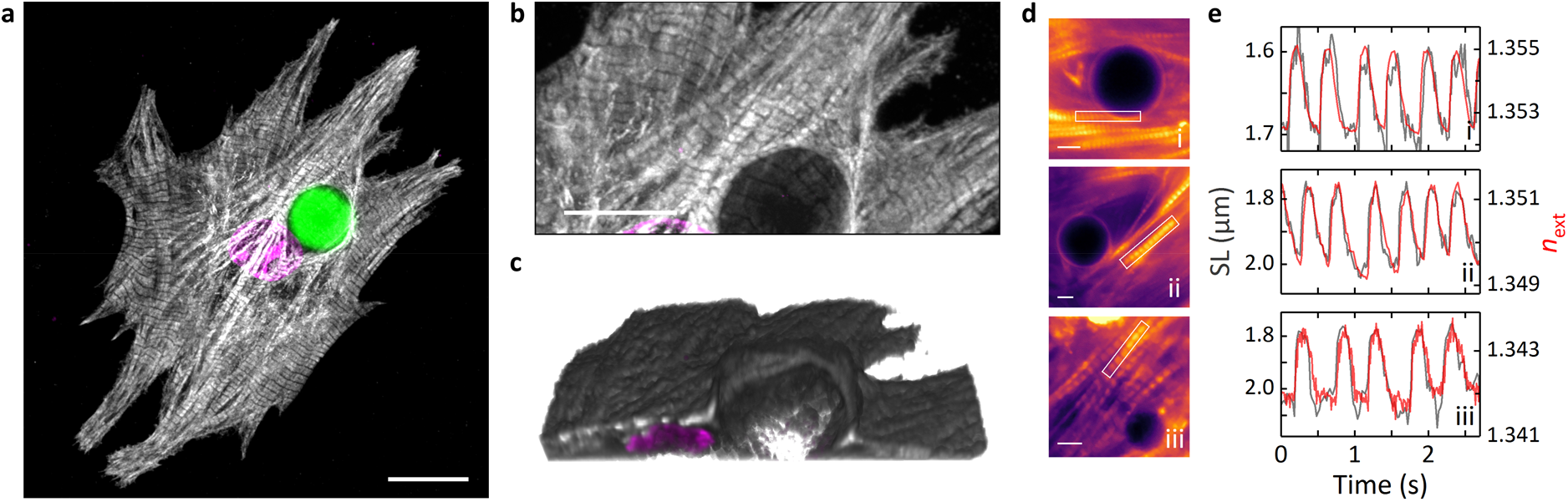
Microlasers monitor cellular contractility. 3D arrangement of myofibrils around microlasers in neonatal cardiomyocytes. **a**, Maximum intensity projection showing the sarcomeric protein cTnT (grey), cell nucleus (magenta) and microlaser (green). Scale bar, 15 μm. **b**, Magnified region around the microlaser and **c**, 3D reconstruction of the same area. The microlaser is omitted to show the arrangement of myofibrils more clearly. Scale bar, 10 μm. **d**, Video rate fluorescence microscopy (Supplementary Video 2) of neonatal mouse CMs with labelled myofibrils. Intracellular microlasers are visible as dark circular objects. Scale bars, 5 μm. **e**, Simultaneously acquired temporal profiles of sarcomere length (SL, grey, left axis, extracted from fluorescence profiles of the myofibrils highlighted by the white rectangles in **d**) and *n*_ext_ (red, right axis, extracted from microlaser spectra). Subfigures labelled according to the images in **d**.

As the microlaser size provides a unique label to identify and track individual cells over time (Supplementary Fig. 6)^13^, we were able to perform repeated monitoring of single neonatal CMs (Fig. 4a). Normalized contractility profiles (Fig. 4b) showed high temporal regularity with minimal beat-to-beat variations in pulse width (FWHM, Fig. 4c) and contraction time (*t*_con_, Fig. 4d). For the example in Fig. 4a, after 42 h, we observe the spontaneous transition into tachycardia which is typically accompanied by increased myocardial tension at elevated beating rates (Bowditch effect), a fundamental process underlying the force-frequency relationship of the heart^21^. At the cellular level, this is caused by increased contractility, which we detected as a step-like increase in the maximum and baseline *n*_ext_ (black arrow in Fig. 4a), allowing simple quantification of relative protein density changes during the entire contraction cycle.

**Fig. 4.**
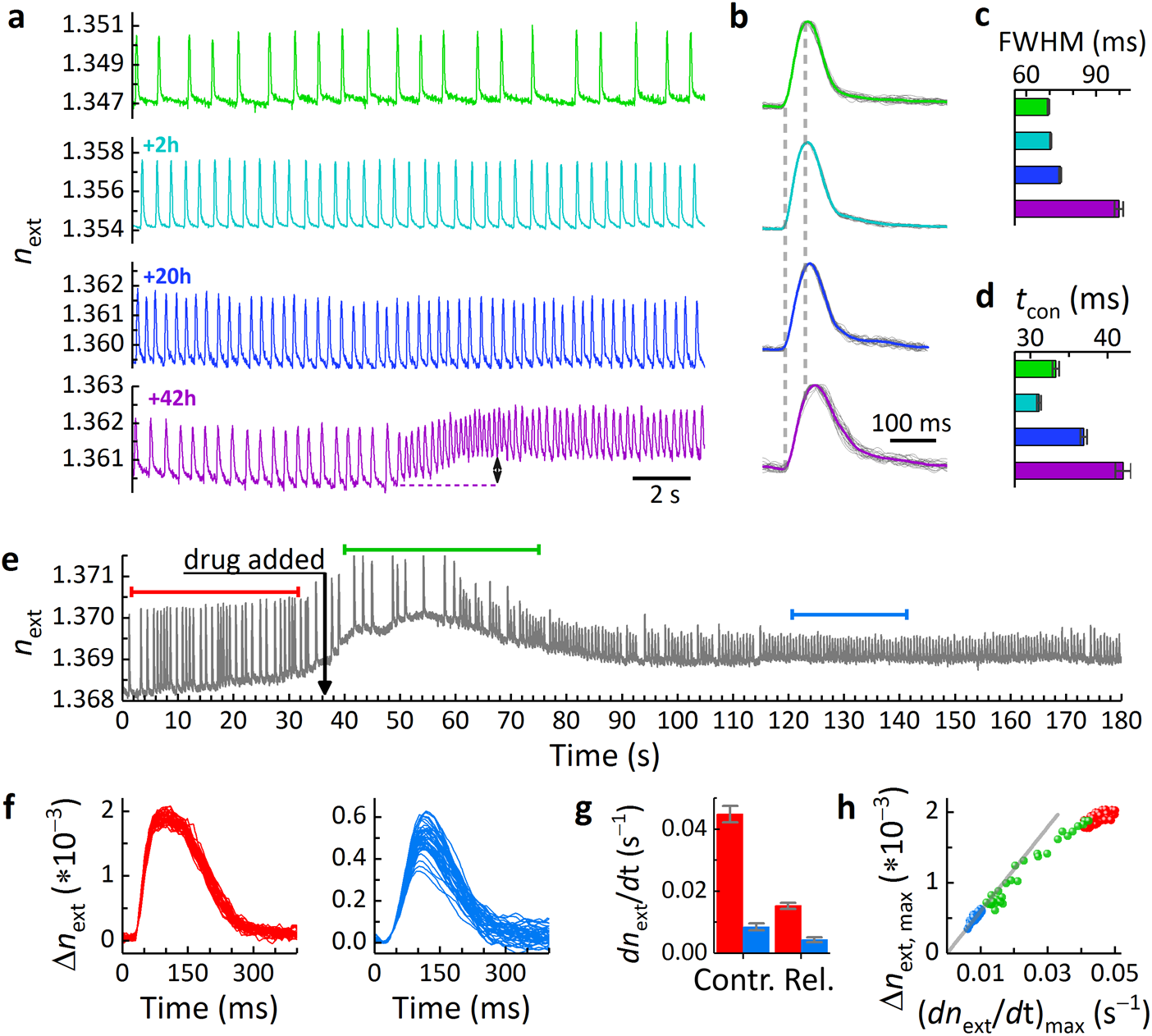
Single cell tracking and contractility sensing under compromised conditions. **a-d**, Microlaser-based tracking and monitoring of a single neonatal CM. **a**, Intracellular *n*_ext_ trace (green) of an individual CM at start of experiment, and characterized again after 2 h (cyan), 20 h (blue) and 42 h (violet). The black arrow marks increased contractility during spontaneous tachycardia. **b**, Normalized *n*_ext_ profiles of traces shown in **a** for n=30-40 cell contractions (grey lines), overlaid by the averaged *n*_ext_ profiles (coloured lines). **c**, Full-width-half-maximum (FWHM) and **d**, average mean contraction time (*t*_con_) of the beating profiles in **b**. **e-h**, Effect of nifedipine on single cell contractility. **e**, *n*_ext_ trace of a spontaneously beating neonatal CM during administration of 500 nM nifedipine (black arrow). **f**, Absolute change in refractive index (Δ*n*_ext_) recorded before (left, red bar in **e**) and after (right, blue bar in **e**) administration of nifedipine. **g**, Average maximum speed of contraction and relaxation for the beating profiles shown in **f**. **h**, Peak refractive index change Δ*n*_ext, max_ plotted as function of the maximum contraction speed with linear fit to nifedipine data. Also shown is the intermediate region (green bar in **e**). Grey line represents linear fit to the data after equilibration of the cell (blue spheres). All error bars represent s.e.m.

Next, we used the quantitative RI transient provided by our laser sensors to assess the effect of the calcium channel blocker nifedipine (Fig. 4e). While the effect of nifedipine on voltage-gated Ca^2+^-channels and subsequent intracellular Ca^2+^ dynamics is well documented^22,23^, the effect on contractility (Fig. 4f) is less well understood as it is difficult to access in neonatal and iPS-derived cardiomyocytes. After administration of nifedipine and following a short period of adaptation, spontaneously beating neonatal CMs showed strongly reduced contraction and relaxation speeds (Fig. 4g), consistent with a reduced concentration of cytosolic Ca^2+^. Furthermore, while we observed that nifedipine increased the pulse-to-pulse variability in Δ*n*_ext_, the time to reach the maximum contraction changed only marginally (Fig. 4f). The lower contraction speed was therefore largely caused by reduced contractility of the cell within the same contraction time which is most likely a result of the calcium dynamics being only slightly affected by nifedipine^22^. Interestingly, this leads to a linear relationship between the mechanical dynamics (contraction speed) and the maximum density change a cell can produce (Δ*n*_ext_), with the latter levelling off with increasing maximum contraction speed (Fig. 4h). Assuming that the saturation contractility of Δ*n*_ext_ = 0.002 observed prior to administration of nifedipine is limited by the number of cross bridges that contribute to force generation, only about 25% of these cross bridges bind to thin filaments in the presence of 500 nM nifedipine.

Microlaser contractility measurements can also be combined with all-optical electrophysiology platforms^22,23^. Simultaneous laser spectroscopy and calcium imaging were performed on fully differentiated mouse CMs that comprise highly organized myofibrils and a transverse tubular system ensuring synchronized calcium release and rapid contraction throughout the cell (Figs. 5a and 5b, Supplementary Video 3). Being non-phagocytic, adult CMs are not able to actively internalise microlasers; so we instead measured spectral changes in the emission of microlasers that were in contact with the cell membrane. Transient profiles of single adult CMs again showed contractions as periodic increases in RI, albeit with smaller amplitude than before (Fig. 5c, Supplementary Fig. 7), demonstrating that Δ*n*_ext_ depends on the volume overlap of the evanescent component of the lasing mode with the myofibrils. However, consistent with our previous observation, the RI transient showed a direct correlation with sarcomere length (Fig. 5c), confirming a contraction-induced change in myofibril density. We also compared the contractility profile to the profile of cytosolic Ca^2+^ and found a characteristic latency time of 30 ms between calcium signalling and force development while the maximum contraction speed coincided with peak Ca^2+^ concentration (Fig. 5c).

**Fig. 5.**
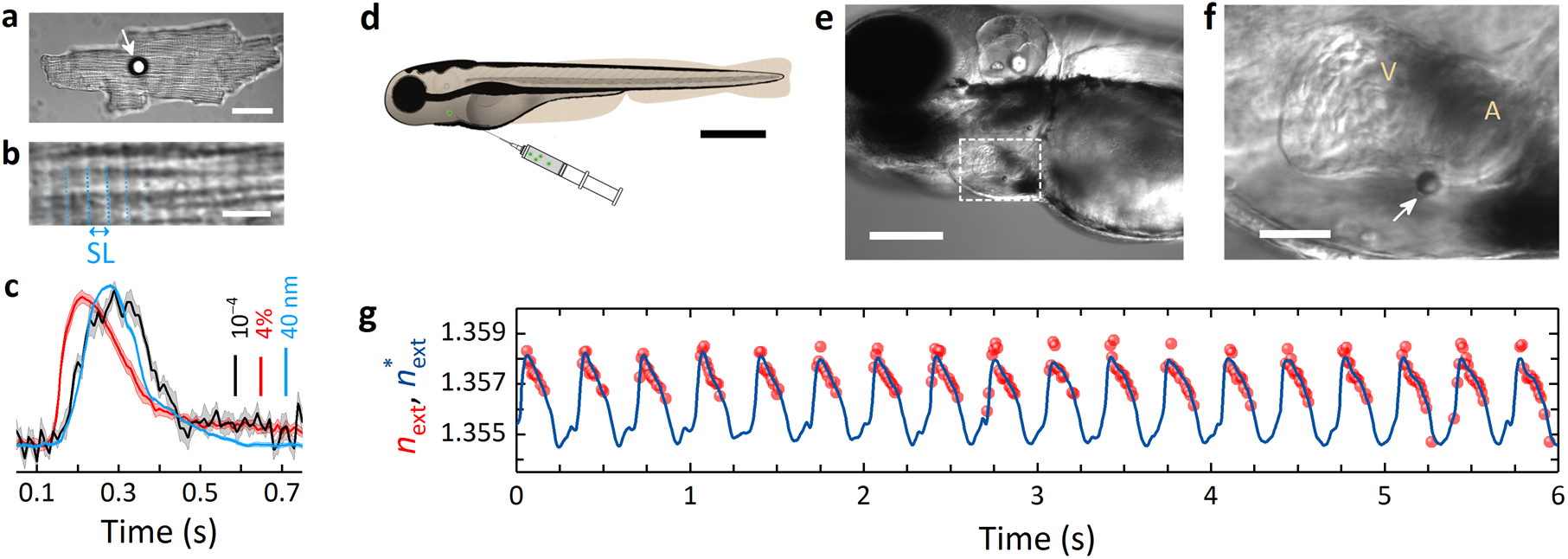
Multimodal sensing and *in vivo* integration. **a**, Extracellular microlaser (white arrow) on top of an adult CM. Scale bar, 30 μm. **b**, Magnified view showing highly organized myofibrils (sarcomere repeat units indicated by dashed blue lines). Scale bar, 4 μm. **c**, Averaged profiles of 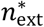 (black), SL (blue), and fluorescent calcium reporter (red). Shaded areas represent s.e.m. of at least 10 contractions. **d-g**, Integration of microlaser into live zebrafish. **d**, Schematic drawing of microlaser injection. Scale bar, 500 μm. **e**, Microlaser attached to the atrium of a zebrafish heart (3 dpf). Scale bar, 200 μm. **f**, Magnified view of the microlaser (arrow). V, ventricle; A, atrium. Scale bar, 50 μm. **g**, *n*_ext_ (red spheres) and 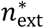 (blue line) calculated using sensitivity calibration.

Having demonstrated intra- and extracellular sensing *in vitro*, we next implemented our technique in live zebrafish, a model organism with remarkable capabilities to repair and regenerate large fractions of the heart^24^. Microlasers were injected by a microneedle (Fig. 5d), placing them at the outer wall of the atrium (Figs. 5e and 5f). Extracellular sensing rather than direct intracellular injection was performed to avoid disruption of the myocardium which at this developmental stage consists of a single layer of cardiomyocytes^25,26^ that is not yet covered by the developing epicardium. Lasing wavelengths again showed the typical red-shift associated with cardiomyocyte contraction (Supplementary Fig. 8). Due to increased tissue scattering and rapid movement (Supplementary Video 4), the intensity of individual modes varied strongly, and the lower intensity TM modes were not detected for a large fraction of the contraction cycle. However, after calibrating the sensitivity of the microlaser from time-points that contained a sufficient number of modes (Supplementary Fig. 9; c.f. Fig. 2d), we were able to construct complete contractility profiles for the beating zebrafish heart (Fig. 5g). A measurement performed at a more posterior position of the atrium revealed a significantly longer systolic plateau (Supplementary Fig. 10), demonstrating locally resolved contractility profiles under *in vivo* conditions.

Restoring cardiac function after severe heart injury remains a major clinical challenge due to the low capacity of the adult mammalian heart to produce new CMs^27^. Current regeneration approaches explore the injection of CMs derived from human embryonic stem cells (hESC) or induced pluripotent stem cells (hiPSC) into the injured heart and the growth of cardiac tissue *in vitro*^3–6^. Multifunctional probes which monitor the long-term integration of injected cells or engineered tissue are urgently needed. Chemical sensing with dye-based or transgenic calcium and voltage reporters are now routinely used for all-optical electrophysiology^25,28^. However, despite their importance, these sensors do not provide insights into the mechanical forces developed by the cells. The processes by which engineered and native cardiac tissue couple mechanically therefore remain unknown^4,29^. Microlaser-based contractility measurements fill this critical gap by monitoring the contractile properties of individual cells during various developmental stages without the need for staining or genetic alteration. Our spectroscopic contractility technique is expected to be more resilient to scattering than imaging-based methods since scattering in biological tissue is elastic and hence does not alter spectral characteristics. Furthermore, the nanosecond-pulsed pumping in combination with single-shot read-out applied here virtually eliminates temporal averaging effects that represent a common source of motion artefacts in intravital confocal or light sheet microscopy.^30^ This can be combined with recent advances in focussing of light deep into scattering tissue^31^, to achieve remote and non-invasive monitoring of cardiac function *in vivo*. By providing single cell specificity, long-term tracking, and reduced sensitivity to scattering, microlasers introduce new possibilities for translational approaches that extend well beyond current microscopy-based techniques, offer reduced complexity, and impose fewer experimental restrictions.

In the future, implementing our recently developed semiconductor WGM nanolasers^32^ or plasmonic nanolasers^33,34^ will improve and simplify internalization further, eliminate any mechanical restriction of the laser probes and drastically reduce the required pump energy. However, surface passivation, heat management and advanced calibration protocols are needed for these single mode lasers before a comparable degree of bio-compatibility and RI sensitivity can be achieved. Furthermore, using high throughput chip-based devices^35^ can enable massively parallel integration of lasers into hiPSC- or hESC-derived cardiomyocytes which in turn would facilitate labelling and monitoring of individual cells from the very early stages of the generation of functional cardiac tissue onwards. Likewise, microlasers can offer functional sensing in newly developed stem cell therapies that are able to restore infarcted tissue^36^.

## Acknowledgements

This research was financially supported by the European Research Council under the European Union’s Horizon 2020 Framework Programme (FP/2014-2020)/ERC Grant Agreement No. 640012 (ABLASE), by EPSRC (EP/P030017/1) and by the RS Macdonald Charitable Trust. S.J.P. acknowledges funding by the Royal Society of Edinburgh (Biomedical Fellowship) and the British Heart Foundation (Grant FS/17/9/32676). M.S. acknowledges funding by the European Commission (Marie Skłodowska-Curie Individual Fellowship, 659213) and the Royal Society (Dorothy Hodgkin Fellowship, DH160102).

## Author contributions

M.S. designed, performed and analysed laser experiments and imaging. L.W. contributed to lasing experiments and B.C. contributed to sarcomere length measurements. I.R.M.B. and L.W. developed refractive index fitting and peak fitting software, respectively. A.M. and M.S. prepared neonatal CM cultures with support from G.B.M. G.B.R. prepared isolated CMs under supervision of S.J.P. S.J.P. and M.S. designed physiological experiments in isolated CMs. C.S.T. supported the preparation of Zebrafish. M.S. and M.C.G. conceived the project and wrote the manuscript with contributions from all authors.

## Competing interests

All authors declare no competing interests.

## Additional information

**Supplementary information** is available for this paper.

## Methods

### Animals

The use of experimental animals was approved by the Animal Ethics Committee of the University of St. Andrews and the University of Edinburgh. The care and sacrifice of animals used conformed to Directive 2010/63/EU of the European Parliament on the protection of animals used for scientific purposes as well as the United Kingdom Animals (Scientific Procedures) Act 1986.

### Cell culture

HL-1 cells were cultured in Claycomb medium (Sigma-Aldrich) supplemented with 100 μM norepinephrine, 10 % (v/v) foetal bovine serum (FBS), 2 mM L-glutamine and 1 % (v/v) penicillin/streptomycin (PS). The cells were stored in T-25 flasks (Fisher Scientific) and incubated at 37°C with 5% CO_2_. Prior to seeding, the flasks were coated with gelatine/fibronectin (0.02% gelatine, 1 mg/ml fibronectin) for at least an hour to improve adherence of the cells. Cells were supplied daily with 1 ml of medium per 3.5 cm^2^ of culture area to maintain and maximise the contractile activity.

### Isolation and culture of neonatal cardiomyocytes

Neonatal mouse hearts were obtained from postnatal day 2 – 3 C57 laboratory mice. Tissue was collected, cleaned and cut into pieces in ice-cold calcium- and magnesium-free Dulbecco’s phosphate buffered saline and digested for 30 min in papain (10 units/ml; Worthington) at 37°C. Treated tissue was dissociated to a single cell suspension by gentle reverse-pipetting in cell culture medium (Dulbecco’s Modified Eagle’s Medium with 25 mM glucose and 2 mM Glutamax, 10 % (v/v) FBS, 1 % (v/v) non-essential amino acids, 1 % (v/v) PS). Non-disaggregated material was allowed to sediment for 2 minutes and the cell suspension pelleted by centrifugation at 200 x g for 5 min. Pelleted cells were resuspended in cell culture medium and pre-plated on an uncoated cell culture flask for 2 – 4 h to enrich cardiomyocytes through surface-attachment of fibroblasts. The cell culture medium containing unattached cells was then recovered from this flask, cardiomyocytes concentrated by centrifugation and seeded at a density of 2 x 10^5^ cells per dish. Prior to seeding, culture dishes (Ibidi) were coated with 0.02 % gelatin/5 μg/ml fibronectin. Cultures were kept in a humidified incubator at 37°C, 5% CO_2_, 95% air. 1 x 10^5^ microlasers (15 μm PS-DVB microspheres stained with Firefli Fluorescent Green, uniformity <12%, Thermo Fisher, UK, 11895052) were added to the dish one day after seeding and incubated over night. Lasing experiments were performed within the next 1-2 days while cultures showed widespread spontaneous contractions for up to two weeks.

### Isolation of adult cardiomyocytes

Adult cardiomyocytes were isolated using an adapted Langendorff-free protocol as previously described.^37^ Isolation solutions used were based on a modified Tyrode’s solution: EDTA buffer (in mM): 5 KCl, 130 NaCl, 0.5 NaH2PO4, 10 HEPES, 10 glucose, 5 Na-pyruvate and 5 ethylenediaminetetraaceticacid (EDTA) titrated to pH 7.8 with NaOH; Perfusion buffer (in mM): 5 KCl, 130 NaCl, 0.5 NaH2PO4, 10 HEPES, 10 glucose, 5 Na-pyruvate and 1 MgCl2 titrated to pH 7.8 with NaOH; Collagenase buffer: 35 mg collagenase type II (Worthington, USA), 50 mg BSA and 15 mg protease (type XIV, Sigma-Aldrich, UK) diluted in 30 ml of perfusion buffer.

Adult C57 mice were killed by cervical dislocation, the chest cavity rapidly opened and descending vessels severed. The right ventricle was injected with 7 ml of EDTA buffer over 1 minute to quickly clear residual blood and stop contraction. The ascending aorta was clamped in situ using haemostatic forceps and the heart excised. The heart was then submerged in EDTA buffer, with a further 10 ml injection of EDTA buffer into the left ventricle over 3 minutes. EDTA buffer was cleared by injection of 3 ml of perfusion buffer into the left ventricle. The heart was then submerged in collagenase buffer, and 30-50 ml of collagenase buffer injected into the left ventricle over 10 minutes. Digestion was taken as complete following a marked reduction in resistance to injection pressure. The digested heart was then transferred to a culture dish containing fresh collagenase buffer and trimmed of any excess non-cardiac tissue. Cardiomyocyte dissociation was completed by gentle trituration using a P1000 pipette. Enzymatic digestion was inhibited by addition of perfusion buffer containing 5 % (v/v) FBS (FBS; Thermo Fisher, UK). Isolated cardiomyocytes were reintroduced to Ca^2+^ by three rounds of 20 minutes sequential gravity settling in perfusion buffer containing 300 μM, 500 μM, and 1 mM CaCl_2_, respectively. Cells were stained (see below) and subsequently transferred into a culture dish (Ibidi) containing 1 mM Ca^2+^ perfusion buffer. After the cells sedimented, 1 x 10^5^ microlasers were added to the dish and lasing experiments were performed within 3 hours of isolation.

### Cardiomyocyte staining

Neonatal cardiomyocytes were labelled with 100 nM SiR-actin overnight. Following isolation, adult cardiomyocytes were loaded with 10 μM X-Rhod-1 AM (λ_ex_= 580 nm, λ_em_= 602 nm; Thermo Fisher, UK) in perfusion buffer containing 1 mM CaCl_2_ for 45 minutes at room temperature. Cells were then washed in 1 mM Ca^2+^ perfusion buffer and left for 15 minutes at room temperature to allow de-esterification of X-Rhod-1 AM.

### Laser spectroscopy

All components for optical pumping and laser spectroscopy were integrated into a standard inverted fluorescence microscope (Nikon, TE2000), equipped with epi fluorescence and differential interference contrast (DIC). The output from a Q-switched and mode-locked diode pumped solid state laser (Alphalas) with wavelength, pulse width and repetition rate of 473 nm, 1.5 ns, and 100 Hz, respectively, was coupled into the objective via a dichroic filter and passed to the sample through either a 60x oil immersion or a 40x long working distance objective. In addition, a further 1.5x magnification was used for sarcomere length tracking. The pump laser was focussed to a 15 μm large spot and a maximum pulse energy of 5-50 nJ was used depending on resonator size and tissue scattering. Emission from the microlaser was collected by the same objective, separated from the pump light by the dichroic and passed to the camera port of the microscope. The image was relayed to a 300 mm spectrometer (Andor) and a cooled sCMOS camera (Hamamatsu) using a series of relay lenses and dichroic beam splitters. The pump laser and spectrometer were synchronized such that each spectrum corresponded to a single pump pulse. During lasing experiments cells were kept in a humidified on-stage incubator system (Bioscience Tools) set to 37°C and purged with 5% CO_2_, 95% air.

Laser thresholds were acquired on the same setup by varying the pump power with a set of neutral density filters. Below threshold, spectra were integrated over 800 pump pulses while above threshold between 100 and 20 pump pulses were used. SNR measurements were performed in a subsequent scan but by integrating over only 1 pump pulse to resemble the conditions of the cardiac measurements. SNR is defined as laser mode peak intensity over fluorescence background.

### Confocal microscopy

Confocal imaging was performed on a Leica TCS SP8 laser scanning microscope with 40× and 63× oil immersion objectives. Neonatal CMs were fixed for 10 min in 4% paraformaldehyde, permeabilized with Triton X-100 and subsequently incubated with the primary cardiac troponin T (cTnT) monoclonal antibody (Thermo Fisher, UK, MA5-12960), the secondary Anti-Mouse IgG CF™ 594 antibody (Sigma-Aldrich, SAB4600092), and DAPI. DAPI, microlasers and myofibrils were excited by sequentially scanned continuous wave lasers with a wavelength of 405 nm, 488 nm, and 594 nm, respectively.

### Multimodal imaging

In addition to the laser coupling spectroscopy optics, a red bandpass filter placed in the dia illumination path of the microscope, a quad-edge epi-luminescence filter cube and additional band pass filters at the spectrograph and camera allowed simultaneous recording of microlaser lasing spectra, and the epi-fluorescence and DIC imaging of cells. Live cell imaging was performed by using an on-stage incubator system.

### Sarcomere length measurements

To determine the average sarcomere length in neonatal mouse cardiomyocytes, myofibrils were fluorescently labelled with SiR-actin (see above) and videos were recorded under epi-illumination conditions at 50 fps using a 60x oil immersion objective (NA 1.4). Raw fluorescence microscopy images were first smoothed by removing statistical noise.^38^ From the smoothed videos intensity profiles were taken along individual myofibrils, covering 5 to 8 sarcomere units. Profiles were then interpolated by a factor of 10, to facilitate an increase in the spatial resolution of the length measurements to about 10 nm that was otherwise limited by the pixel size of the camera and magnification of the microscope. Interpolated profiles were smoothed using the Savitzky-Golay method. Minima in the intensity profiles were tracked through time at 20 ms intervals. Once the separation between the first and last minima was determined, it was divided by the number of sarcomeres to calculate the average sarcomere length in each frame. In adult cardiomyocytes, sarcomere length measurements were performed using DIC videos recorded at 100 fps by using the ImageJ plugin SarcOptiM.^39^ Briefly, a fast Fourier transformation algorithm is used to extract the regular spacing in a line profile plotted along the longitudinal axis of the cell. Adult cardiomyocytes were electrically paced at 1 Hz with Platinum wire bath electrodes by applying 8 ms square voltage pulses with a maximum electric field of 30 V/cm.

### *In vivo* zebrafish experiments

All zebrafish embryos used in our experiments were under the age of 5 days post fertilisation (dpf). Embryos were collected from random matings and then correctly developmentally staged. Fertilised eggs were transferred at the 2–8 cell stage to 10 cm culture dishes at 28.5°C with systems water replaced every 24 h. When necessary, larvae were anaesthetised with MS-222 (tricaine methanesulfonate, 40 μg/ml, Sigma-Aldrich). Microlasers were injected into the sinus venosus region of 3 dpf embryos with a micropipette (pulled on a Sutter P97) attached to a Narishige IM-300 microinjector, whilst viewed on a stage of a Leica M16F stereo microscope. Lasing experiments were performed at room temperature.

## Data availability

All research data presented in this study will be made available on the University of St Andrews online depository PURE.

## Code availability

The custom-made computer code is available from the corresponding authors upon request.

## Notes

#### Summary of Updates

Supplementary Information and Video 1-4 have been added.

