## Supplementary Information for "Monitoring contractility in single cardiomyocytes and whole hearts with bio-integrated microlasers"

**Quantification of microlaser uptake**

Internalization of spherical microlasers was quantified following a previously developed uptake assay.[1] Briefly, microlasers were coated with biotin and incubated with cardiac cells for 24 h, after which point cells were fixed. To label extracellular microlaser, the cell impermeable streptavidin-Atto 647N conjugate was added. Intracellular lasers are characterized by showing only green bulk fluorescence of the laser but negligible red surface fluorescence (Supplementary Fig. 1). Automated analysis of epi-fluorescence images is used to obtain quantitative uptake efficiencies (Supplementary Fig. 2).

**WGM resonances and refractive index sensing**

Light that circulates in a homogeneous dielectric sphere is strongly concentrated by total internal reflection and positive interference. The discrete angular resonance frequencies $\omega_n$ of the supported WGMs can be linked to the resonance size parameter $x_n^l$:

$$\omega_n = 2\pi c \lambda_0^{-1} = \frac{x_n^{(l)} c}{n_{ext} \cdot r} \ , \qquad \text{(Eq. 1)}$$

where $c$ is the speed of light in vacuum, $\lambda_0$ the vacuum wavelength, and $r$ the radius of the sphere. $x_n^{(l)}$ can be accurately calculated by an asymptotic series and explicitly depends on the azimuthal and radial mode numbers $n$ and $l$, respectively, the polarization, as well as on the refractive index of the sphere ($n_s$) and the surrounding ($n_{ext}$):[2]

$$x_n^{(l)}(n, l, n_s, n_{ext}) = \frac{v}{m} - \frac{\zeta_l}{m}\left(\frac{v}{2}\right)^{\frac{1}{3}} + \sum_{k=0}^{k_{max}} \frac{d_k(m, \zeta_l)}{v^{k/3}(m^2-1)^{(k+1)/2}} \ . \qquad \text{(Eq. 2)}$$

Here, the refractive index contrast is defined as $m = n_s/n_{ext}$, $\zeta_l$ denotes the $l$th zero of the Airy function, $v = n + 1/2$, and $d_k(m, \zeta_l)$ are the polarization dependent coefficients of the asymptotic expansion which have been calculated up to $k_{max} = 8$.[2] For small polystyrene spheres ($n_s$=1.6) used in this study, only first order radial modes ($l = 1$) are observed,[1] while azimuthal mode numbers typically range from $n =$ 100-200.

The resonance frequencies of dielectric spheres (and in turn lasing wavelengths of our WGM microlasers, which are referred to as mode positions in the following) are strongly dependent on $n_{ext}$ and can therefore be used for refractive index sensing. A look-up table approach is applied to extract the refractive index in the surrounding of the sphere ($n_{ext}$) together with the sphere radius $r$ from each measured WGM lasing spectrum. Firstly, precise experimental WGM positions are obtained from the centre position of a Gaussian peak fit to the respective spectrum. With a spectral resolution of the spectrometer system used in this study of 55 pm, the mode position after peak fitting can be determined with about 1 pm accuracy. We then calculate the difference between the measured resonance positions and a large database of calculated spectra were $n_{res}$ and $r$ are systematically varied using a resolution of $\Delta n_{res} = 0.00002$ and $\Delta r = 100$ pm. Typically, about 5 million spectra are calculated, each comprised of all TE and TM modes within a spectral range of 500 to 540 nm (which corresponds to the spectral region over which the fluorescent dye in our microlasers provides significant net optical gain). To reduce the influence of the greater uncertainty in the experimental mode position of low intensity resonances (fitting noise), the size of the resonator was averaged before reiterating the time series data by keeping the radius fixed and sweeping only over the refractive index value.

**Sensitivity of WGM microlasers, detection limit and single mode calibration**

The refractive index change induced by a beating cardiomyocyte can be utilised to calibrate the refractive index sensitivity $S$ of the microlaser, allowing calculation of the linearly approximated refractive index ($n^*_{\mathrm{ext}}$) from

$$n^*_{\mathrm{ext}} = S * \lambda_{\mathrm{TE/TM}} + \mathrm{n}_0 \quad . \qquad \text{(Eq. 3)}$$

Unfortunately, one cannot obtain a general value for $S$ as it depends strongly on the resonator size as well as on several (microscopic) parameters like dye concentration, surface roughness, and the refractive index contrast between cell and microlaser. However, for small perturbations and for individual lasers, the resonance wavelength is expected to change linearly with refractive index. Thus, the contraction induced changes in cellular refractive index provide an ideal perturbation to calibrate the sensitivity of the microlaser. Supplementary Fig. 3 summarizes the procedure to obtain the sensitivity and compares the results obtained using Eq. 3 to the full optical modelling and look-up table approach described above. For resonators with a diameter between 13 and 17 µm, we find that $S$ typically ranges between 0.055-0.065 $nm^{-1}$. It should be noted that each mode has a different sensitivity and that we generally calibrated the most intense mode.

**Effect of microlaser on myofibrillar network**

The organization of myofibrils around intracellular microlasers was investigated by confocal laser scanning microscopy (Supplementary Fig. 5). Cells with internalized microlasers maintain a network of myofibrils, predominantly located in a thin layer but significantly extending in the region of the microlaser. Myofibrils are found to stretch along the bottom, side and eventually up to the top of the microlasers as shown in the 3D reconstructions. Another characteristic feature is the close distance of cell nucleus and microlaser which are either in direct contact or separated by only a few micrometer in the centre of the cell.

**Origin of increased systolic refractive indices**

The relative increase of the external refractive index ($\Delta n_{\mathrm{ext}}$) related to the contraction of cardiomyocytes can be linked to an increase in protein concentration ($\Delta c_{\mathrm{p}}$) inside the evanescent volume of the microlasers, providing a method for dynamic measurements of local protein concentration (or protein density, which is an equivalent measure).

Refractive index and protein concentration are directly connected via the well documented mean protein refractive index increment ($dn/dc$) of 0.190 ml/g,[3] a parameter that is only slightly dependent on the molecular weight and exact chemical composition of proteins. It is often used to link refractometric measurements of proteins to their concentration in solution. In our experiments, the maximum experimentally observed increase in refractive index is $\Delta n_{\mathrm{ext,\,max}} = 0.003$ (Fig. 2d, main text), which corresponds to a maximum observed increase in protein concentration of:

$$\Delta c_{\mathrm{p,\,max}} = \Delta n_{\mathrm{ext,\,max}} \times \left(\frac{dn}{dc}\right)^{-1} = (0.003/0.19)\,\frac{\mathrm{g}}{\mathrm{ml}} = 0.0158\,\frac{\mathrm{g}}{\mathrm{ml}}$$

The total protein concentration of vertebrate skeletal muscle fibrils has been determined to $c_{\mathrm{fibril}} = 0.185\ \mathrm{g/ml}$,[4] comparable to the protein concentration of 0.2 g/ml typically found in mammalian cells.[5] The relative increase in protein concentration is therefore $\frac{\Delta c_{\mathrm{p,\,max}}}{c_{\mathrm{fibril}}} = \frac{0.0158}{0.185} * 100\% = 8.5\%$. For the 3 examples of correlative measurements of sarcomere length and refractive index change (Fig. 3b, main text) a maximum increase in protein concentration of 7.7% (i), 6.5% (ii) and 5.4% (iii) is calculated. Assuming that the total number of proteins and thus their mass remains unchanged during the contraction cycle implies that the myofibril volume decreases by about 5-8% as protein concentration scales inversely with volume.

We can now compare these values to measurements of the sarcomere unit cell volume of intact rat cardiomyocytes. By combining X-ray scattering and laser diffraction measurements, *Yagi et al.* observed that the lattice volume decreases linearly with decreasing sarcomere length.[6] Similar changes were observed by others but dismissed in favour of a constant volume interpretation.[7,8] Using the linear relationship provided by *Yagi et al.*, a change from 2 µm at diastole to 1.8 µm at systole (compare to Fig. 3b, main text) would result in a decreased lattice volume of 6.4%. Our results are therefore in good agreement with direct structural measurements on myofibrils and support the findings that myofibrils undergo a significant change in lattice volume upon cardiomyocyte contraction. The increased systolic concentration of proteins increases the local refractive index and is the origin of the red shift of lasing modes that we observe consistently throughout our experiments.

**Cell tracking with intracellular lasers**

WGM microlasers have been proposed as size-encoded single cell tags where the microlaser diameter provides a characteristic parameter that distinguishes one resonator from the other. The idea is based on the high accuracy by which the size of the resonator can be determined form the multimode lasing spectrum and further requires a high geometrical uniformity (sphericity) of the microlaser. We performed cell tracking experiments by repeatedly measuring intracellular microlasers over the course of 2 days (Fig. 3d, main text). The diameter of the microlaser is calculated by independently extracting the microlaser size from the first 10 spectra of a time series (Supplementary Fig. 6). These 10 diameters are then averaged and the mean diameter is subsequently used to evaluate the intracellular refractive index ($n_{ext}$) for the whole time series. Repeated measurements of the same microlaser typically yield size variations of a few nanometers, significantly larger than the measurement uncertainty. This is attributed to slight deviation of the microlasers from perfect sphericity which must be considered when estimating the maximum number of cell tags. The polymer microspheres used in this study have a size range from 10 to 20 µm. Setting size increments of 5 nm, 2000 cells can be uniquely distinguished consistent with previous estimates.[9]

1. Schubert, M. *et al.* Lasing in Live Mitotic and Non-Phagocytic Cells by Efficient Delivery of Microresonators. *Sci. Rep.* **7,** 40877 (2017).
2. Schiller, S. Asymptotic expansion of morphological resonance frequencies in Mie scattering. *Appl. Opt.* **32,** 2181 (1993).
3. Zhao, H., Brown, P. H. & Schuck, P. On the Distribution of Protein Refractive Index Increments. *Biophys. J.* **100,** 2309–2317 (2011).
4. Reedy, M. K. & Lucaveche, C. in 29–45 (Springer, Boston, MA, 1984). doi:10.1007/978-1-4684-4703-3_4
5. Milo, R. What is the total number of protein molecules per cell volume? A call to rethink some published values. *BioEssays* **35,** 1050–1055 (2013).
6. Yagi, N. *et al.* Sarcomere-length dependence of lattice volume and radial mass transfer of myosin cross-bridges in rat papillary muscle. *Pflugers Arch. Eur. J. Physiol.* **448,** 153–160 (2004).
7. Irving, T. C., Konhilas, J., Perry, D., Fischetti, R. & de Tombe, P. P. Myofilament lattice spacing as a function of sarcomere length in isolated rat myocardium. *Am. J. Physiol. Circ. Physiol.* **279,** H2568–H2573 (2000).
8. Farman, G. P., Allen, E. J., Gore, D., Irving, T. C. & de Tombe, P. P. Interfilament Spacing Is Preserved during Sarcomere Length Isometric Contractions in Rat Cardiac Trabeculae. *Biophys. J.* **92,** L73–L75 (2007).
9. Humar, M., Upadhya, A. & Yun, S. H. Spectral reading of optical resonance-encoded cells in microfluidics. *Lab Chip* **17,** 2777–2784 (2017).

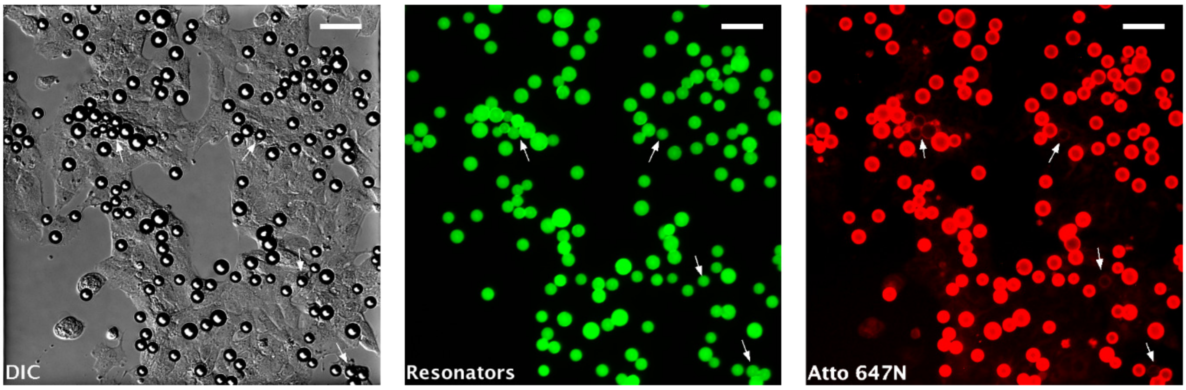


**Supplementary Figure 1 | Quantification of microlaser uptake in the cardiac cell line HL-1.** The uptake assay consists of a series of DIC (left), green fluorescence (center) and red fluorescence (right) images. Microlasers without red surface staining (white arrows) are counted as intracellular. Scale bars, 50 µm.

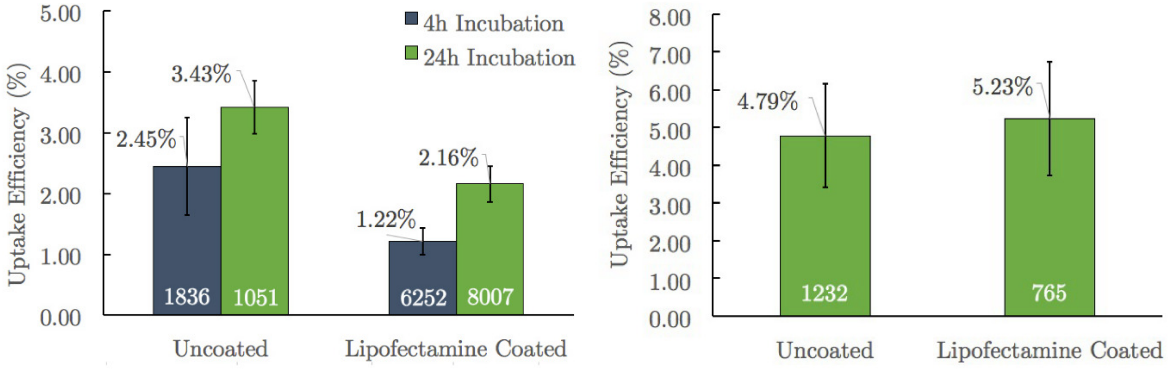


**Supplementary Figure 2 | Quantifying microlaser internalization.** Uptake efficiency of HL-1 (left) and neonatal cardiomyocytes (right). Lipofectamine coating, which has been shown to increase uptake efficiency in other cell types[1], has no positive effect, consistent with the reported low liposomal transfection efficiency of cardiac cells. Numbers on top and bottom of error bars denote mean uptake efficiency and number of analysed resonators, respectively. Error bars represent standard error of the mean.

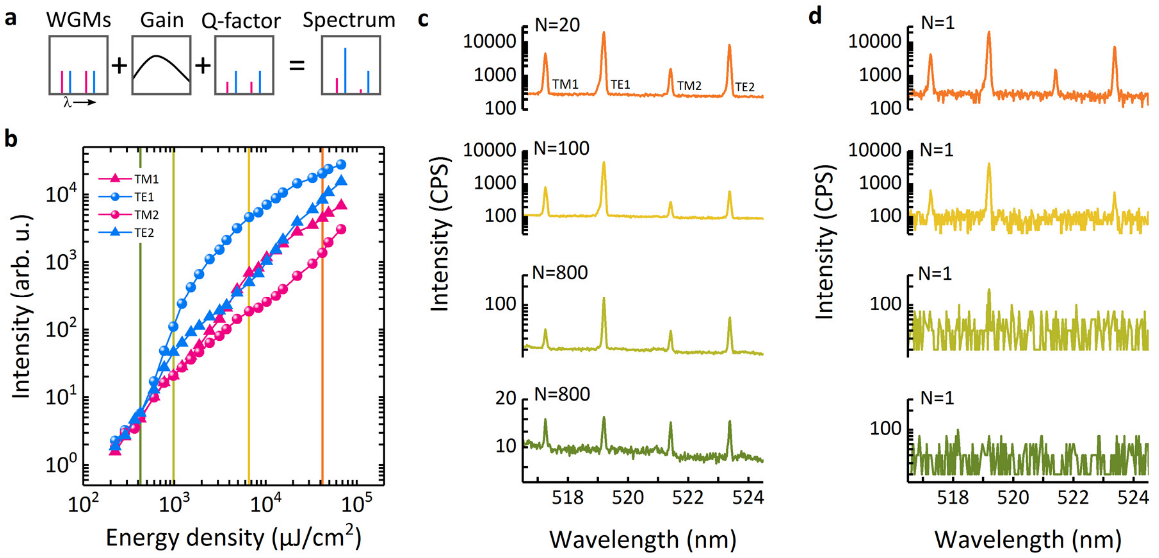


**Supplementary Figure 3 | Threshold analysis of multimode microlasers. a**, Schematic of the different factors contributing to the characteristic emission profile of lasing modes in spherical microlasers. Starting with the geometrically defined modes, convolution with the gain spectrum and the Q-factor of individual modes result in a mode profile that at high pump power is dominated by a single TE mode. **b**, Measured threshold characteristics of 4 WGMs. The dominating TE mode has the lowest threshold and quickly increases in intensity over the fluorescence background. Other modes subsequently start lasing at significantly higher thresholds. With increasing pump power, mode competition reduces the relative intensity difference between the modes, causing a saturation in the signal-to-noise ratio (see Fig. 2e, main text). Example spectra at selected pump energy densities (corresponding to the coloured vertical lines in **b**) **c**, averaged over *N* pump pulses or **d**, with single pulse excitation as applied during CM measurements. Under single pulse excitation a minimum pump energy density of 1000 μJ/cm$^2$ is required to detect the dominating lasing mode while the least intense TM mode (TM2) can only be detected once it passes the lasing threshold. The spectra and threshold characteristics shown here are reproducible for each microlaser and representative for microlasers in the investigated size range of 10 to 20 μm.

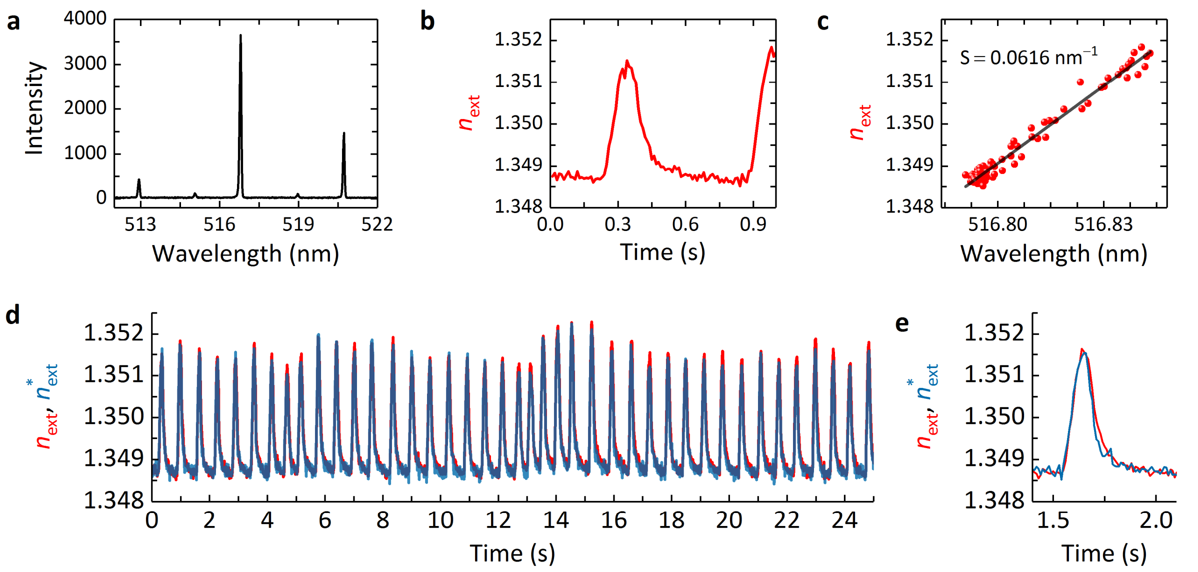


**Supplementary Figure 4 | Protocol to determine the refractive index sensitivity $S$ of microlasers. a**, Microlaser spectrum containing 5 WGMs. **b**, Calculated $n_{ext}$ for a 1 s time interval of a typical contraction profile. **c**, For all time points, the calculated $n_{ext}$ is plotted over the spectral position of the mode that is to be calibrated. Linear fit (black line) to the data ($R^2$=97.8%) from which $S$ and $n_0$ are derived. **d**, Plot of the linearly approximated refractive index ($n^*_{ext}$) for the complete 25 s measurement. The refractive index obtained by using the mode position of all 5 WGMs and the look-up table approach ($n_{ext}$) is shown for comparison. **e**, Detailed comparison of $n^*_{ext}$ and $n_{ext}$ for a single contraction. The dynamics and amplitude of the contraction are well resembled by the linearly approximated signal while the slightly lower noise in the $n_{ext}$ trace is due to an effective averaging of the 5 modes compared using data from just a single mode.

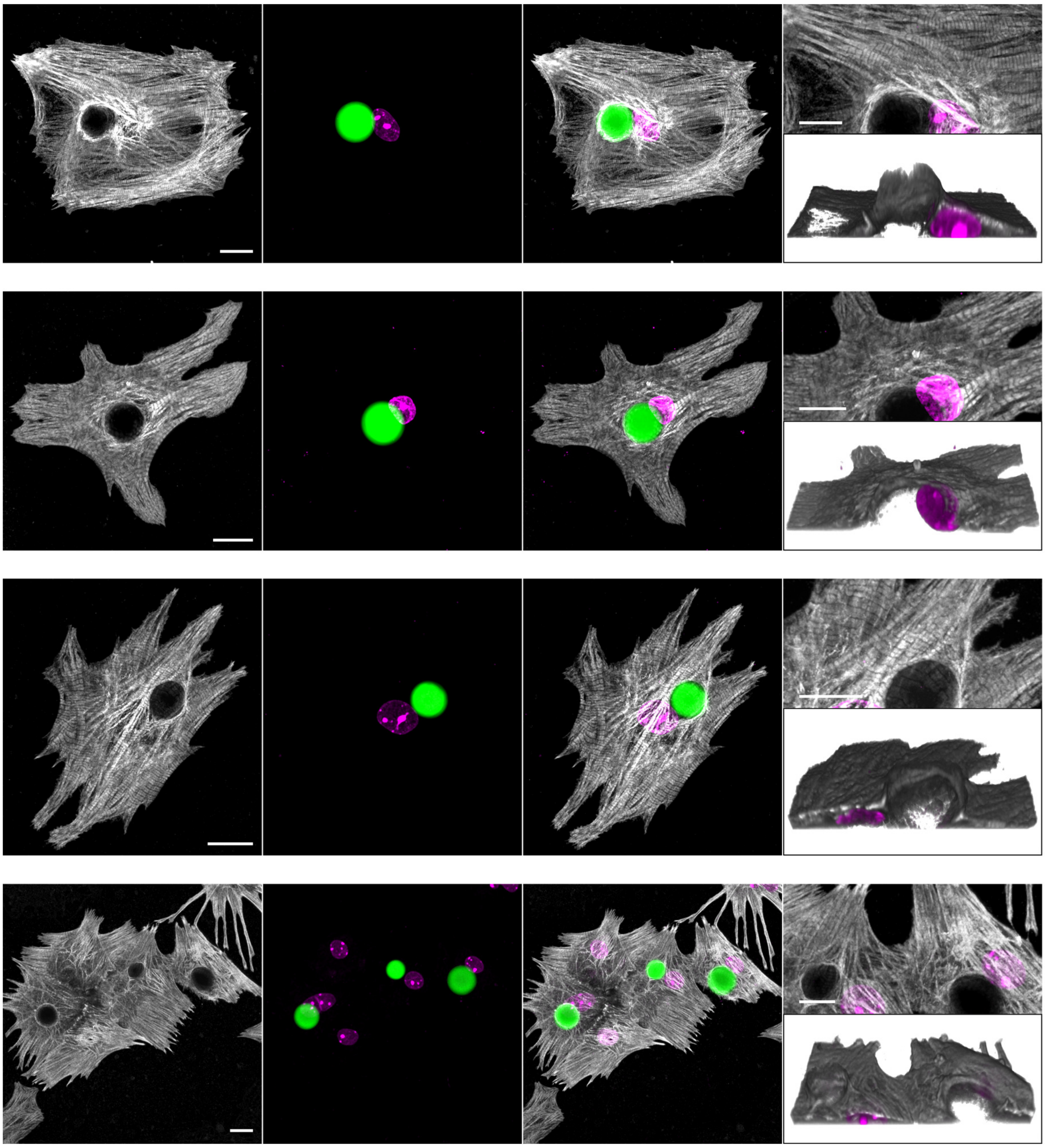

**Supplementary Figure 5 | Myofibrils organize around intracellular microlasers.** Confocal microscopy images (maximum intensity projection) showing the sarcomeric protein cTnT (grey), microlasers (green) and cell nucleus (magenta). Scale bars, 15 µm. Microlasers are predominantly located at the cell centre, often forming a direct contact with the cell nucleus. Furthermore, myofibrils stretch around the microlaser and cover a large fraction of the surface by projecting fibrils out of the otherwise flat morphology of the cells. This is shown in more detail in the magnified regions around the microlaser and the 3D reconstructions of the same area. The microlaser has been omitted from these images to visualize the 3D arrangements of myofibrils. Scale bars, 10µm.

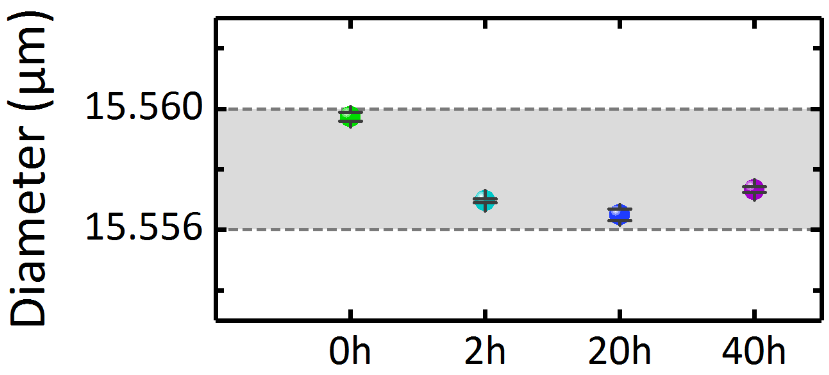


**Supplementary Figure 6 | Change of apparent microlaser diameter during an extended experiment.** Mean microlaser diameter obtained by averaging 10 WGM spectra at each time point. Error bars display the standard error of the mean which is smaller than 100 pm for all time points. The grey area indicates the 4 nm range within which the calculated size of microlasers typically varies.

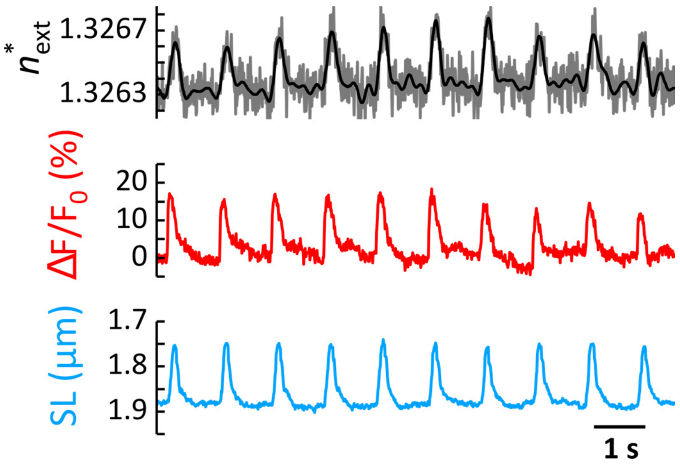


**Supplementary Figure 7 | Transient profiles of an adult CM electrically paced at 1 Hz**. Top: $n^*_{\text{ext}}$ recorded by the extracellular microlaser (Black=smoothed, grey=unprocessed). Middle: relative change in fluorescence $\Delta F/F_0$from Calcium reporter XRhod1. Bottom: sarcomere length (SL) extracted from Fourier analysis of DIC microscopy videos.

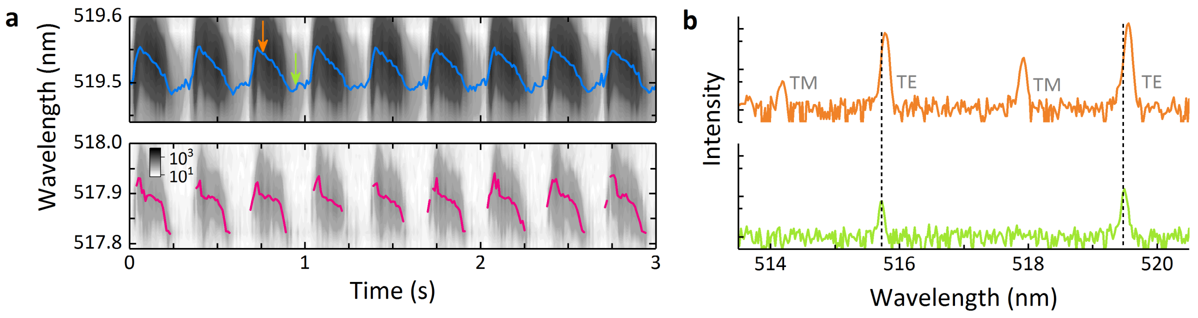


**Supplementary Figure 8 | Details of WGM spectra measured inside live zebrafish.** **a**, Fitted mode position of a pair of TE (blue) and TM (pink) WGMs (Fig. 4, main text; Video 4). **b**, WGM spectra recorded at systole (orange) and diastole (green) showing variations in the peak intensity due to fluctuations in pump and collection efficiency upon movement of the resonator. Vertical dashed lines indicate TE mode positions in diastole.

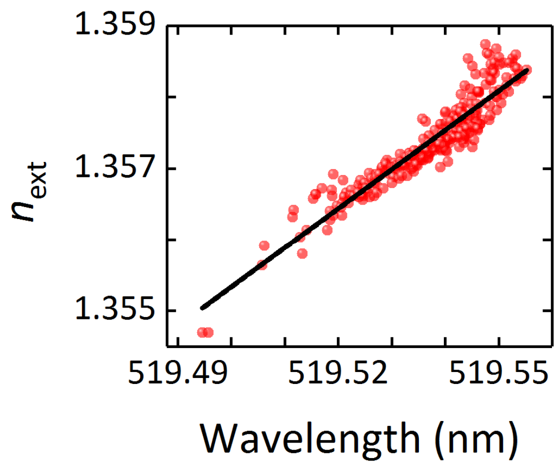


**Supplementary Figure 9 | Microlaser calibration inside live zebrafish.** Refractive index sensitivity $S$ obtained by a linear fit to $n_{ext}$ (obtained from spectra that contain at least 4 WGM, see Supplementary Fig. 8) versus the mode position of the most intense WGM ($S$ =0.055 nm$^{-1}$, $R^2$=87.7%).

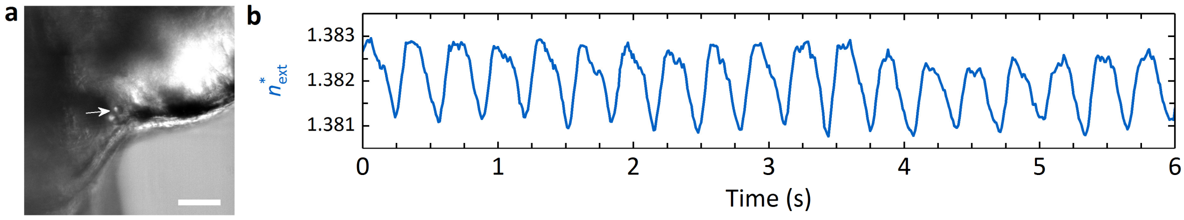


**Supplementary Figure 10 | Additional in vivo measurements.** Measurement of a microlaser taken at a more posterior position compared to the one shown in Fig. 4 (main text).

**Video 1** Visualisation of the data acquisition and analysis workflow. **a**, DIC microscopy time-lapse video of a spontaneously beating neonatal cardiomyocyte. The internalized microlaser is seen as circular object in the centre of the cell**.** Images were acquired with a 60x oil immersion objective at a rate of 100 Hz. **b**, Whispering-gallery mode spectrum of the microlaser acquired simultaneously with DIC images. Dotted vertical lines mark the laser mode position at diastole. **c**, Centre position of the laser mode at around 519.75 nm obtained after peak fitting of the spectrum in **a**. **d**, External refractive index $n_{\mathrm{ext}}$ calculated by the look-up table algorithm that, at each time point, compares the positions of the 5 laser modes shown in **b** to a database of simulated spectra. All panels are synchronized in time and played in real time. Experiments repeated in more than 8 independent repeats for a total of N>80 cells.

**Video 2** Fluorescence microscopy time-lapse video of the 3 neonatal cardiomyocytes shown in Fig. 3a (main text). Cells were labelled with SiR-actin to visualize sarcomeric actin inside the myofibrils. The microlasers are seen as dark circular objects. Experiment performed in triplicate for a total of N=12 cells.

**Video 3** Multimodal imaging of cellular calcium dynamics and contractility. **a**, Fluorescence microscopy time-lapse video of the adult cardiomyocyte shown in Fig. 4a (main text). The cell was labelled with the calcium-sensitive dye XRhod1 and images were recorded at a rate of 50 Hz. **b**, Smoothed external refractive index $n^{*}_{\mathrm{ext}}$ measured by a microlaser located on top of the cell. A longer time trace is shown in Supplementary Fig. 7. **c**, Fluorescence intensity profile normalized to the intensity at resting phase obtained from the video shown in **a**. Experiments performed in duplicate for a total of N=5 cells. All panels are synchronized in time.

**Video 4** *In vivo* contractility measurement by a microlaser attached to the atrium of a zebrafish embryo (3 dpf). The simultaneously acquired microlaser spectra and the calculated external refractive index are shown in Supplementary Fig. 8 and Fig. 4g (main Text), respectively. A measurement performed at a more posterior position of the heart is shown in Supplementary Fig. 10.
